# Hydrogenotrophic methanogenesis and distinct microbial assemblages fostered by dauciform roots of *Cladium jamaicense*

**DOI:** 10.1101/2025.10.06.680723

**Authors:** Kevin R. Montiel, John S. Kominoski, Anna K. Simonsen

## Abstract

Nutrient availability regulates ecosystem processes, and plant roots strongly influence nutrient cycling through microbial interaction in the rhizosphere. In the extremely phosphorus-limited Florida Everglades, *Cladium jamaicense* (sawgrass) develops specialized roots, called dauciform roots, which release carboxylates that mobilize soil-bound nutrients. Although methanogenesis is a well-documented process in wetlands, the composition of methanogenic communities across separate root-associated compartments is not as well characterized.

We investigated microbial community composition and predicted functional pathways across bulk soil, the general rhizosphere soil, and the rhizosphere of dauciform roots in calcareous (marl) soils near Everglades National Park. Functional predictions were inferred from 16SrRNA gene data using PICRUSt2 with taxonomic assignments referenced against SILVA v138.2 using rANOMALY.

Alpha and beta community analyses revealed significant differences among compartments. Dauciform roots harbored the lowest Shannon diversity, whereas bulk soils supported the most distinct assemblages. Microbial communities clustered strongly by compartments, with compartment identity explaining 66% of the variation (p = 0.001). Pairwise comparisons showed the strongest separation between bulk and dauciform soils. Furthermore, functional predictions showed enrichment of hydrogenotrophic methanogenesis sequences in dauciform roots, while acetoclastic methanogenesis was most abundant in rhizosphere soils further emphasizing their distinct communities.

Our preliminary results demonstrate that root-associated compartments foster distinct microbial assemblages with implications for key ecosystem processes, including methanogenesis. These findings highlight how root traits in oligotrophic systems influence carbon cycling and potential methane pathways, contributing to broader insights into microbial community assembly and ecosystem processes in nutrient-limited wetlands.

**Highlights:**

- First genomic investigation of the dauciform root rhizosphere.
- Root & dauciform rhizosphere harbor microbial communities distinct from bulk soil.
- Predicted enrichment of hydrogenotrophic methanogens in dauciform roots.

---

Nutrient availability is a primary determinant of ecosystem processes, and plants influence nutrient cycling through interactions with soil microbes through their roots (Grayston et al., 1998). Root systems can alter the biological and chemical conditions of soils. By releasing exudates in the form of labile carbon and carboxylates, roots stimulate microbial activity and increase nutrient availability in the surrounding soil region, known as the rhizosphere (Oburger et al., 2009). In nutrient-limited wetlands such as the ultra-oligotrophic Florida Everglades, phosphorus scarcity shape plants root system, drive underground processes, and affect the distribution of microbial communities (Noe, 2001). Thriving in water saturated soils, methanogens contribute to the production of methane, a greenhouse gas that’s at least twenty-five times more potent than CO_2_ (Aliyev et al., 2020). Methanogenesis in wetlands is well documented, yet most studies emphasize soil processes (Bridgham et al., 2013), with comparatively few examining the rhizosphere where roots traits influence microbial community structures. However, far less is known about how methanogenic communities are structured across separate root-associated compartments, where variation in root architecture and carbon inputs may foster distinct microbial assemblages.

The Everglades offers a natural setting to study these dynamics, where the dominant native macrophyte, *Cladium jamaicense* (sawgrass), contributes to the structure and function of the soil environment (Larsen et al., 2010). Depending on nutrient availability, *C. jamaicense* can produce specialized lateral roots, known as dauciform roots (Richards & Olivas, 2019), which are dense bundles of root-hairs that increase root surface area and enhance nutrient acquisition (Fig. 1) (Shane et al., 2006). As exudation hotspots, dauciform roots help mitigate phosphorus limitations by releasing carboxylates that mobilize soil-bound nutrients (Playstead et al., 2006); however, their influence on rhizosphere microbial communities remains understudied. We present the first preliminary evidence that the root compartments of *C. jamaicense* harbor distinct microbial communities with differences that extend to the composition of predicted methanogenesis assemblages. We further provide evidence that dauciform roots influence microbial composition, create unique microbial niches, and harbor distinct assemblages that differ from both the general rhizosphere and the non-root-associated bulk soil.

**Figure 1.**
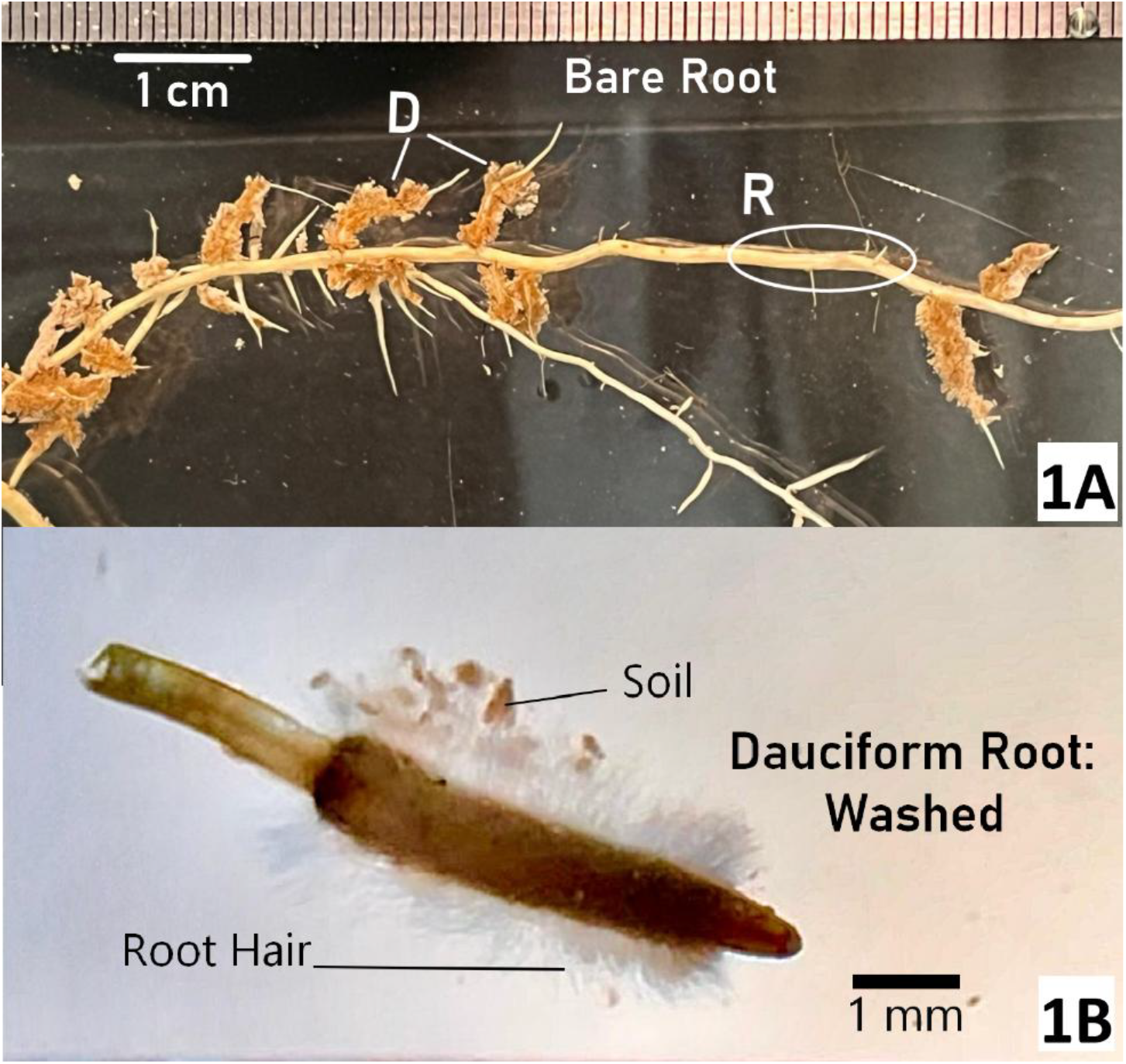
Root morphology of *Cladium jamaicense*. 1A: Bare root with distinct clusters of dauciform roots (D), displaying their distinctive carrot-like shape along the general rhizosphere regions (R). 1B: Washed dauciform root displaying dense root hairs and trapping soil particles, illustrating their specialized morphology for nutrient acquisition. Scale bars: 1A = 1 cm, 1B = 1mm

To assess differences in microbial community structure among *C. jamaicense* compartments, we compared microbial communities of root-associated soil and bulk soil, the portion not directly influenced by roots. A one-meter quadrat was placed in an oligotrophic, calcareous soil (marl), sawgrass-dominated wetland along the eastern boundary of Everglades National Park (25º24’N, 80º33’W). Four mature plants and their surrounding soil were excavated (n=4). The surrounding soil was collected as bulk soil, and the plants were then shaken to remove any soil not directly adhering to the roots. Soil adhering to fine absorptive roots, corresponding to first-to-third order roots, was scraped off and collected as rhizosphere soil (McCormack et al., 2015). Dauciform roots were separated and vortexed in sterile tubes to dislodge soil tightly bound to their root hair clusters.

Following manufacturer’s instruction, DNA extraction was done using DNeasy PowerSoil Kit (QIAGEN). The NC State Genomic Sciences Laboratory performed 16SrRNA gene amplification using the primer pair 341F-805R, which amplifies the V3–V4 region, and performed on a NextSeq 2000 (Illumina) using a NextSeq 300 PE P1 flow cell. We processed reads with rANOMALY (Theil & Rifa, 2021), which integrates DADA2 for denoising and resolving amplicon sequence variants (ASV) and subsequently assigned taxonomy against SILVA v.138.2 (Quast et al., 2013). Community analyses were conducted in Phyloseq with visualizations in rANOMALY. Functional predictions based off 16SrRNA (PICRUSt2; Douglas et al., 2020) were compared in STAMP (Parks et al., 2014) using ANOVA with Tukey-Kramer post hoc tests, and p-values were corrected by the Benjamini-Hochberg FDR method.

Alpha and beta community analyses show significant differences in microbial communities across soil compartments. Shannon diversity index differed significantly by soil type (Fig. 2A; p = 0.036), with the dauciform soils being the lowest, bulk soils values were tightly clustered while the rhizosphere soil compartment averaged slightly higher accompanied by greater variability. Principal coordinates analysis ordination (PCoA), based on Bray-Curtis dissimilarities, revealed clear clustering by soil compartment. Points represent individual soil samples, and together the soil compartments explained 66% of the variation in community structure according to PERMANOVA (Fig. 2D: R^2^ = 0.66, p = 0.001). Pairwise PERMANOVA confirmed significant differences among all compartments, with the strongest separation between bulk soil and dauciform roots (R^2^ = 0.62, p = 0.038), followed by bulk versus the rhizosphere (R^2^ = 0.55, p = 0.038). Together, these results indicate that bulk soils harbor the most distinct assemblages, whereas root-associated compartments also differed significantly (R^2^ = 0.43, p = 0.038) but showed partial overlap, indicating greater similarity to each other compared to bulk soil. These patterns may indicate that bulk soil functions as a reservoir of microbial diversity, whereas dauciforms foster specialized assemblages and distinct communities in the general rhizosphere.

**Figure 2.**
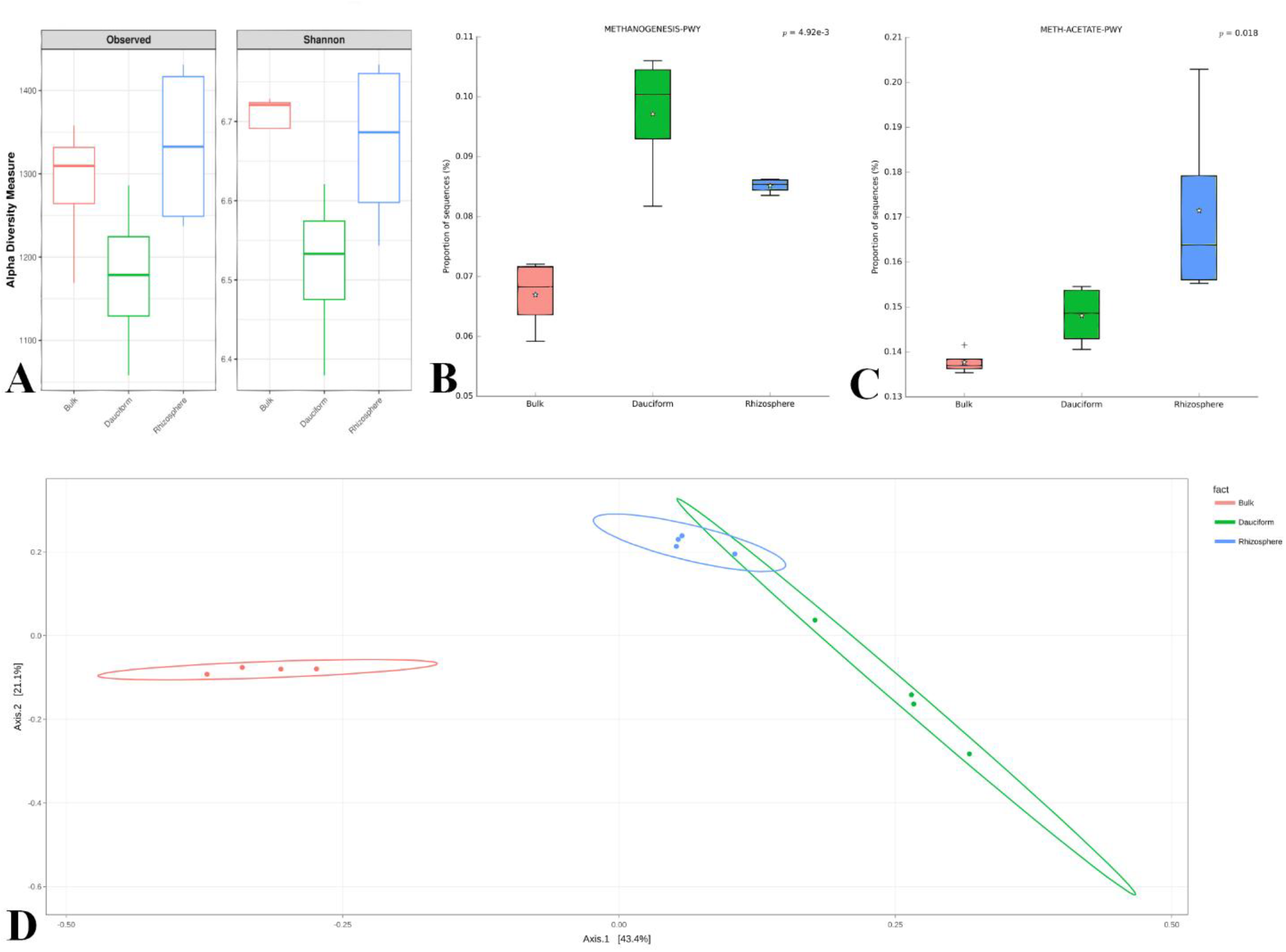
Community structure and functional potential for methanogenesis across soil compartments. **2A**. Observed richness did not differ significantly among compartments. In contrast Shannon diversity was significant (*), indicating that dauciform roots harbored lower diversity compared to bulk and rhizosphere samples. **2B**. Hydrogenotrophic methanogenesis pathway is significantly enriched (**) in dauciform root samples. **2C**. Predicted acetate-dependent methanogenesis pathways show enriched abundance in rhizosphere soils (*), reflecting a potential functional distinction of microbial communities across root-compartments. **2D**. PCoA ordination reveals distinct clustering by compartment, supported by PERMANOVA (**), with bulk soil forming the most distinct assemblages. Methanogenesis pathways were generated using PICRUSt2. Statistical comparisons were performed in rANOMALY and STAMP. Asterisks denote significance: p < 0.05 (**\***), p < 0.01 (**\*\***).

We applied PICRUSt2 and STAMP to assess the distribution of classified sequences assigned to methanogenesis pathways across the soil compartments. The proportion of predicted sequences associated with the hydrogenotrophic methanogenesis (from H_2_ and CO_2_) pathway was most abundant in dauciform roots, followed by the general rhizosphere and the lowest in the bulk soil (Fig. 2B: p = 4.92 × 10^−3^). In contrast, the proportion of predicted acetoclastic methanogenesis pathway sequences, methane formation from the breakdown of acetate, were enriched in the general rhizosphere, with lower enrichment in the dauciforms and the bulk compartments (Fig. 2C: p = 0.04). Notably, the relative proportion of acetoclastic methanogenesis sequences was double that hydrogenotrophic sequences, consistent with reports indicating that methanogenesis through acetate predominates in wetlands (Aliyev, 2020; Bridgham et al., 2013; Conrad, 1999).

Our preliminary findings indicate that root-associated compartments not only contain distinct microbial communities but also suggest *C. jamaicense* roots create favorable micro-environments that support unique methanogens. Carbon inputs from *C. jamaicense* roots, coupled with root strategies that increase nutrient availability, may establish localized hotspots for methanogenesis. These processes are reflected in distinct, inferred hydrogenotrophic and acetoclastic methanogenic pathways.

The greater relative abundance of predicted hydrogenotrophic compared to acetoclastic methanogenesis sequences in dauciform roots underscores how specialized root traits influence carbon cycling pathways in oligotrophic wetlands. Further experimental work is needed to disentangle and determine how root traits directly mediate the filtering of microbial communities and shape methane production, contributing to the broader question of how organisms persist and adapt in ultra-low nutrient wetland systems like the Florida Everglades.

## Data Availability Statement

The data that support the findings of this study are openly available in **Figshare** at https://doi.org/10.6084/m9.figshare.30282565.v1, reference number 30282565.

## Acknowledgements

This material was developed in collaboration with the Florida Coastal Everglades Long-Term Ecological Research (FCE-LTER) program under National Science Foundation Grant No. DEB-2025954. We are grateful for the Kominoski Lab from FIU for their in-kind support in granting access to the field site, which made this research possible.

